# A novel mechanism of gland formation in zebrafish involving transdifferentiation of renal epithelial cells and live cell extrusion

**DOI:** 10.1101/353524

**Authors:** Richard W. Naylor, Alan J. Davidson

## Abstract

Transdifferentiation is the poorly understood phenomenon whereby a terminally differentiated cell acquires a completely new identity. Here, we describe a rare example of a naturally occurring transdifferentiation in zebrafish in which kidney distal tubule epithelial cells are converted into an endocrine gland known as the Corpuscles of Stannius (CS). We find that this process requires Notch signalling and is associated with the cytoplasmic sequestration of the Hnf1b transcription factor, a master-regulator of renal tubule fate. A deficiency in the Irx3b transcription factor results in ectopic transdifferentiation of distal tubule cells to a CS identity but in a Notch-dependent fashion. Using live-cell imaging we show that CS cells undergo apical constriction *en masse* and are then extruded from the tubule to form a distinct organ. This system provides a valuable new model to understand the molecular and morphological basis of transdifferentiation and will advance efforts to exploit this rare phenomenon therapeutically.

## Introduction

Transdifferentiation is the poorly understood process by which a mature cell is converted to a completely different cell type (Tosh & Slack, 2002). Under normal biological settings, transdifferentiation is a rare phenomenon, although it can be readily induced experimentally by forcing the expression of genes that direct cell fate changes, such as the classic example of the MyoD transcription factor being able to convert fibroblasts into muscle lineage cells (Tapscott et al., 1988). The process of transdifferentiation can be either direct, whereby a mature cell converts seamlessly into another mature cell type, or indirect, whereby there is a requirement for dedifferentiation to a more immature intermediary cell type, followed by differentiation into the new fate (Merrell & Stanger, 2016; Thowfeequ, Myatt, & Tosh, 2007). Examples of transdifferentiation occurring in a developmental setting are uncommon, the best studied cases are in *C. elegans* embryos with the indirect transdifferentiation of rectal epithelial Y cells into cholinergic motor neurons (Jarriault, Schwab, & Greenwald, 2008) and the formation of MCM interneurons from AMso glial cells (Sammut et al., 2015).

In vertebrates, direct transdifferentiation is largely limited to the adult setting where it is associated with response to injury. For example, ablation of pancreatic β-cells induces the transdifferentiation of resident α-cells to β-cells in both mice and zebrafish (Thorel et al., 2010; Ye, Robertson, Hesselson, Stainier, & Anderson, 2015). Similarly, in the liver, chronic injury promotes the conversion of hepatocytes to biliary epithelial cells through the combined action of the Notch and Hippo signalling pathways (Yanger et al., 2013). Cases of indirect transdifferentiation in vertebrates include the well known example of lens regeneration in amphibians following lentectomy (Stone, 1967), in which retinal pigmented epithelial cells initiate expression of pluripotency genes (Maki et al., 2009), dedifferentiate and then mature into lens cells (Sanchez Alvarado & Tsonis, 2006). Indirect transdifferentiation is considered to occur in some cancers, via the epithelial-to-mesenchymal transition and dedifferentiation that often accompanies tumourigenesis (Fang et al., 2005; Maddodi & Setaluri, 2010; Maniotis et al., 1999; Shekhani, Jayanthy, Maddodi, & Setaluri, 2013). In summary, while transdifferentiation *in vivo* is possible under normal and pathogenic settings, it remains a rare and poorly understood phenomenon.

The zebrafish offers a visually accessible vertebrate model with which to study cell fate changes in the context of organogenesis. The embryonic kidney (pronephros) is particularly well suited for these studies because of its readily visualised position within the embryo and a high degree of understanding of how cell division, differentiation and morphogenesis are coordinated during organ formation (I. A. Drummond et al., 1998; Majumdar, Lun, Brand, & Drummond, 2000; Naylor & Davidson, 2016; Naylor, Przepiorski, Ren, Yu, & Davidson, 2013; Naylor et al., 2016; R. A. Wingert & Davidson, 2008, 2011; Rebecca A Wingert et al., 2007). The zebrafish pronephros is analogous to the filtering units in the mammalian kidney (nephrons) and consists of a midline-fused blood filter (glomerulus), attached to bilateral renal tubules that extend to the cloaca (I. A. Drummond & Davidson, 2010; I. A. Drummond et al., 1998; R. A. Wingert & Davidson, 2008; Rebecca A Wingert et al., 2007). The tubules are subdivided into functionally distinct segments consisting of the proximal convoluted tubule (PCT), the proximal straight tubule (PST), the distal early tubule (DE), and the distal late segment (DL; (Fig.1 and Rebecca A Wingert et al., 2007)). Each tubule segment expresses a specific set of genes that defines its functional differentiation. The PCT and PST are associated with bulk re-absorption of solutes from the filtrate and express a wide variety of solute transporters (Blaine, Chonchol, & Levi, 2015; Ullrich & Murer, 1982; Rebecca A Wingert et al., 2007). In contrast, the DE and DL segments express fewer transporters, suggesting that they function more to fine-tune the composition of the filtrate. For example, functionality of the DE segment is conferred by the expression of *slc12a1*. encoding a Na-K-Cl co-transporter (Rebecca A Wingert et al., 2007). The transcription factor Hnf1b is a major regulator of renal tubule identity in both zebrafish and mammals (Heliot et al., 2013; Massa et al., 2013; Naylor et al., 2013). In *hnf1b*-deficient zebrafish embryos, solute transporter expression fails to initiate in the tubule segments and the kidney tubules remain as a simple, non-transporting, tubular epithelium (Naylor et al., 2013).

**Figure 1:**
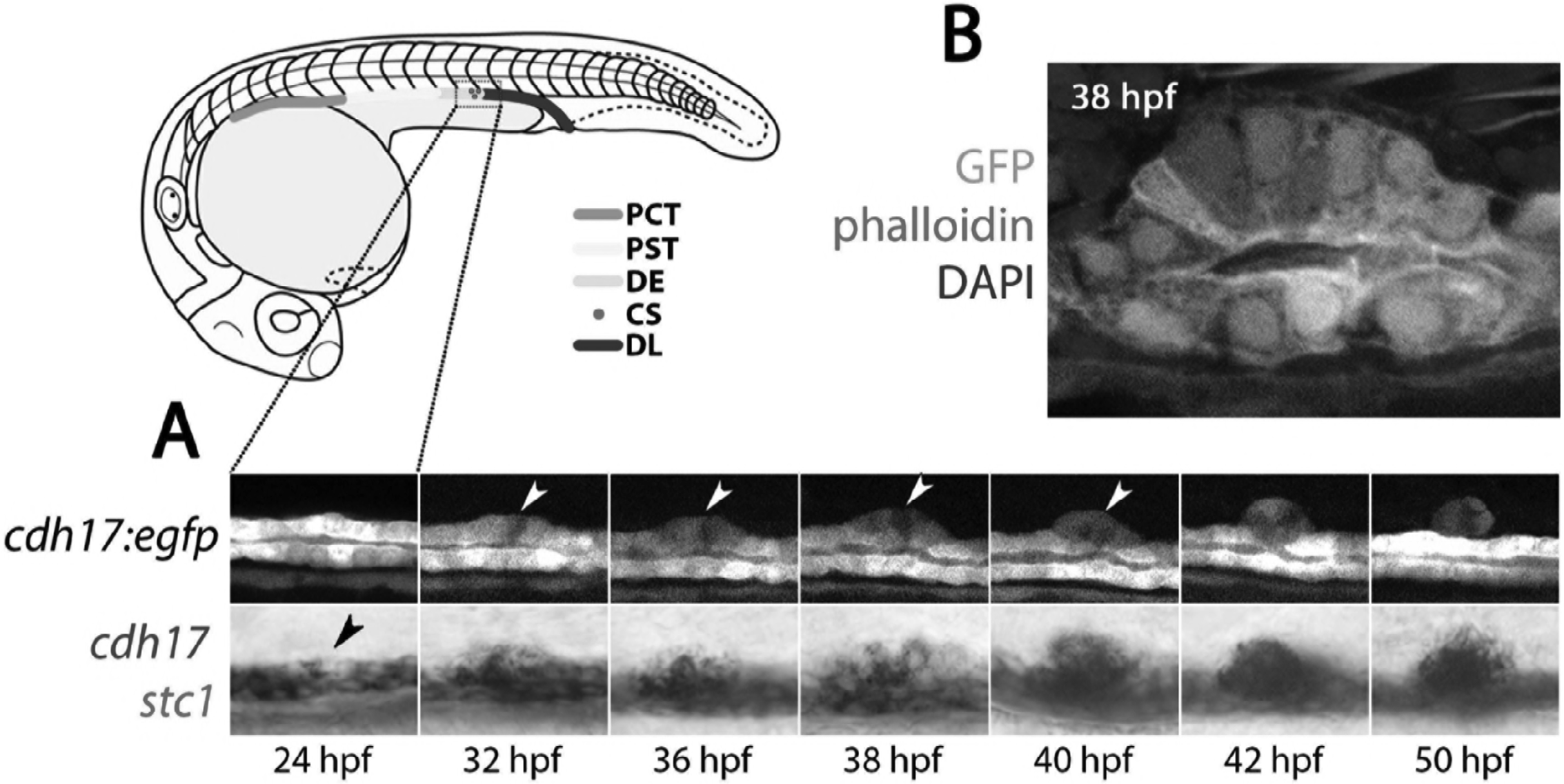
CS extrusion is achieved by apical extrusion of multiple cells. A) Lateral views of a stage series analysis of CS extrusion in a single live *Tg(cdh17:egfp)* embryo (top panels) and embryos fixed at the stages shown and stained for *cdh17*(purple)/ stc1(red). B) Image of a sagittal section through the pronephros of a *Tg(cdh17:egfp)* embryo co-labelled with Phalloidin (F-actin, red) and DAPI (nuclear stain, blue) at the site of the extruding CS at 38 hpf. Abbreviations: PT, Proximal tubule; DE, Distal early tubule segment; CS, Corpuscles of Stannius; DL, Distal Late tubule segment; hpf, hours post fertilisation.

Above the pronephric tubules, and initially forming at the junction between the DE and DL segments, is positioned an endocrine gland called the Corpuscles of Stannius (CS, Fig.1). The CS gland in teleost fish is responsible for secreting Stanniocalcin-1 (Stc1), a hormone involved in the homeostatic regulation of calcium (Cheng & Wingert, 2015; Tseng et al., 2009). The CS glands consist of lobes of epithelial cells that are separated by strands of connective tissue containing blood vessels and nerve tracts, which is quite distinct from the cuboidal polarised epithelium that constitutes the pronephric tubule (Cohen, Pang, & Clark, 1975; Menke, Spitsbergen, Wolterbeek, & Woutersen, 2011). As such, the CS gland is a functionally and structurally distinct organ that shares little homology, other than an epithelial state, to the kidney. Despite this, early observations have led to the suggestion that the CS gland originates from the pronephric tubules, although its formation has not been extensively characterised.

In this study, we investigate the origin of the CS gland and discover that it arises by a previously uncharacterised process of direct transdifferentiation from DE tubule cells followed by apical constriction and extrusion. We find that the molecular pathways that control this transdifferentiation event involve Notch signalling, cytoplasmic sequestration of the renal identity factor Hnf1b, and inhibitory signals conferred by the Iroquois transcription factor Irx3b. Together, these results demonstrate a rare example of direct transdifferentiation under normal physiological conditions in a vertebrate.

## Results

### CS cells originate in the renal tubule and are extruded to form a gland

To better understand CS gland formation we first examined expression of *stanniocalcin-1* (*stc1*), a marker of CS fate, and *cdh17*, a pan-renal tubule marker that encodes a cadherin involved in cell-to-cell adhesion (Horsfield et al., 2002). We found that at 24 hours post-fertilisation (hpf), transcripts for *cdh17* are down-regulated in the posteriormost portion of the DE segment (Fig. 1A). Concomitant with this, the first *stc1*^+^ cells appear in this region and increase in number. By 32 hpf the *stc1*^+^ cells appear as a prominent bulge on the dorsal side of the tubule and at 50 hpf, they are found as discrete, and separate, structures on top of each pronephric tubule (Fig. 1A). These observations suggest that CS cells arise from renal epithelial cells via transdifferentiation and are then physically extruded from the tubule.

To visualise the process of CS gland formation, live time-lapse imaging was performed on *Tg(cdh17:egfp)* embryos from 24 to 50 hpf (Fig.1A and Supplementary Movie 1). Presumptive CS cells were observed to bulge out of the dorsal wall of the tubule concomitant with the constriction of their apical membranes. Immunostaining of sagittal cross-sections through the CS region at this stage showed that the cells adopt a wedge shape with narrowed apical membranes (marked by Phalloidin^+^ F-actin, Fig.1B). As apical constriction increased, we observed that the adjacent non-CS epithelial cells moved closer together and the CS cells protruded dorsally as an arch two cell layers wide (Fig.2A). The epithelial cells on either side of the CS cells eventually met around 42 hpf, concomitant with the extrusion of the CS gland from the tubule (Fig. 1A and Supplementary Movie 1). We found that during extrusion from the tubule, the *epithelial cell adhesion molecule* gene (*epcam*) remains continuously expressed in CS cells (Fig.2B) and the major epithelial cell adhesion protein Cadherin-1 (Cdh1) is retained on the basolateral membranes, suggesting that the epithelial status of CS cells is not changed during extrusion (Fig.2A). After CS cell extrusion, Cdh1 can be detected at the interface between CS and tubular cells and is prominent on the lateral membranes of CS cells, similar to renal epithelial cells (Fig.2A). However by 50 hpf, Cdh1 immunostaining is largely lost from the interface between CS cells and tubular cells but remains on the basal surface of some CS cells (Fig.2A). In addition, F-actin staining, which demarcates the apical surface of tubular cells, is lost in the extruded CS cells (Fig.2A). Taken together, these results support a model in which renal epithelial cells transdifferentiate directly into CS cells, without a loss of epithelial character, extrude from the tubule following apical constriction, and form a distinct ball of unpolarised glandular epithelium.

**Figure 2:**
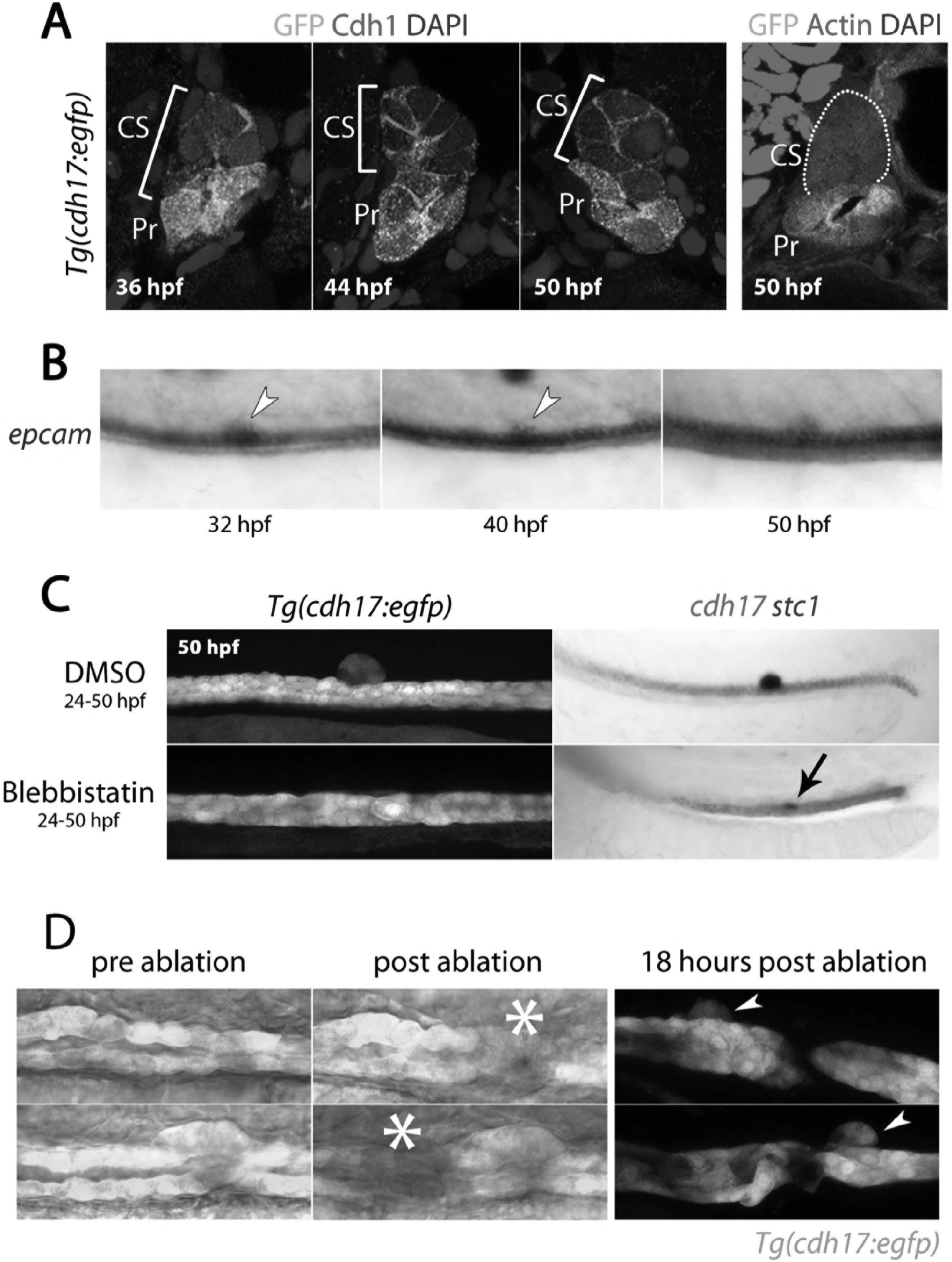
DE to CS transdifferentiation is direct and extrusion is mediated by apical constriction. A) Panels show transverse sections through the CS gland of *Tg(cdh17:egfp)* embryos at the stages indicated. Green fluorescence is from the endogenous GFP, Cdh1 is labelled red and nuclei are labelled blue (DAPI). B) Panels show lateral views of an extruding CS gland in embryos at the indicated stages labelled with *epcam.* C) Panels lateral views of the trunk that show *Tg(cdh17:egfp)* embryos (left) and *cdh17* (red)/ *stc1* (purple) double staining (right) in control and Blebbistatin treated embryos. Black arrow indicates the position of *stc1*^+^ cells in the Blebbistatin treated embryo. D) Panels show brightfield/ fluorescence in *Tg(cdh17:egfp)* embryos before and after laser ablation (site of ablation indicated with asterisk). Panels on the right are of embryos 18 hours post ablation and arrowheads highlight the extruded CS gland.

### Role of apical constriction, proliferation and neighbouring cells during CS gland extrusion

To determine if CS cell extrusion was dependent upon apical constriction, embryos from 24 hpf onwards were incubated in Blebbistatin (10 μM), an inhibitor of Myosin IIA that blocks the contraction of the apical ring of F-actin filaments, which is essential for apical constriction in other contexts (Kovacs, Toth, Hetenyi, Malnasi-Csizmadia, & Sellers, 2004; Lewis, 1947; Sawyer et al., 2010). We found that the CS cells failed to extrude in all Blebbistatin-treated embryos (n=57) and instead, *stc1*^+^ cells remained within the tubule (Fig.2C). Thus we conclude that apical constriction of CS cells is a necessary prerequisite for extrusion from the renal tubule.

We next examined the role of cell proliferation in CS gland formation by performing EdU incorporation experiments. This analysis showed that CS cells only proliferate once they have been extruded from the tubule (n=6 at each time-point, Fig.S1A). Inhibition of proliferation between 20 hpf (20-somite stage) and 26 hpf with hydroxyurea and aphidicolin did not reduce the initial number of *stc1*^+^ cells that arise in the tubule nor does it block CS cell extrusion (n=27/27, Fig.S1B). Consistent with CS cells undergoing cell division following their extrusion, fewer *stc1*^+^ cells were observed in Blebbistatin-treated embryos at later stages compared to controls, suggesting that there are two phases of CS cell formation: an intra-tubule transdifferentiation phase where a small number of CS cells are generated and a post-extrusion expansion phase.

Previously studied examples of live cell extrusion describe an actomyosin contractile ring that forms in cells neighbouring the extruding cell (Eisenhoffer et al., 2012; Gu, Forostyan, Sabbadini, & Rosenblatt, 2011). As all pronephric tubule cells contain apically localised F-actin, it was not possible to determine if an equivalent actomyosin ring forms in the cells surrounding the CS cells. However, laser ablation to sever the tubule (and any interconnected contractile filaments) immediately anterior or posterior to the forming CS gland did not prevent CS extrusion (n=6, Fig.2D), suggesting that neighbouring cells do not play an active role in CS extrusion.

### Nuclear export of Hnf1b occurs in DE cells that transdifferentiate to CS

We next investigated the molecular mechanisms governing the transdifferentiation of DE tubule cells into CS cells. We first examined Hnf1b as this transcription factor is critical for establishing and maintaining renal epithelial cell fate but is not expressed in the extruded CS gland ((Naylor et al., 2013 and Fig.S2)). Immunostaining of embryos at 18 hpf showed that all cells in the renal tubule display nuclear localised Hnf1b (Fig.3A). However from 22 hpf onwards, we found that Hnf1b is lost from the nucleus in presumptive CS cells and becomes localised to the apical cytoplasm in a speckled pattern (Fig.3A, see Fig.S3 for lateral view). As this cytoplasmic localisation of Hnf1b precedes the onset of *stc1* expression it suggests that loss of Hnf1b activity (via cytoplasmic sequestration) may be an early event in the transdifferentiation of renal to CS fate. To provide further support for this notion we examined *hnf1b*-deficient embryos as we reasoned there might be ectopic CS cell formation in the absence of Hnf1b activity. In line with this, we detected a broad stretch of *stc1* expression in the middle of the pronephric tubule, largely corresponding to the region that would normally form the DE segment (Fig.3B).

**Figure 3:**
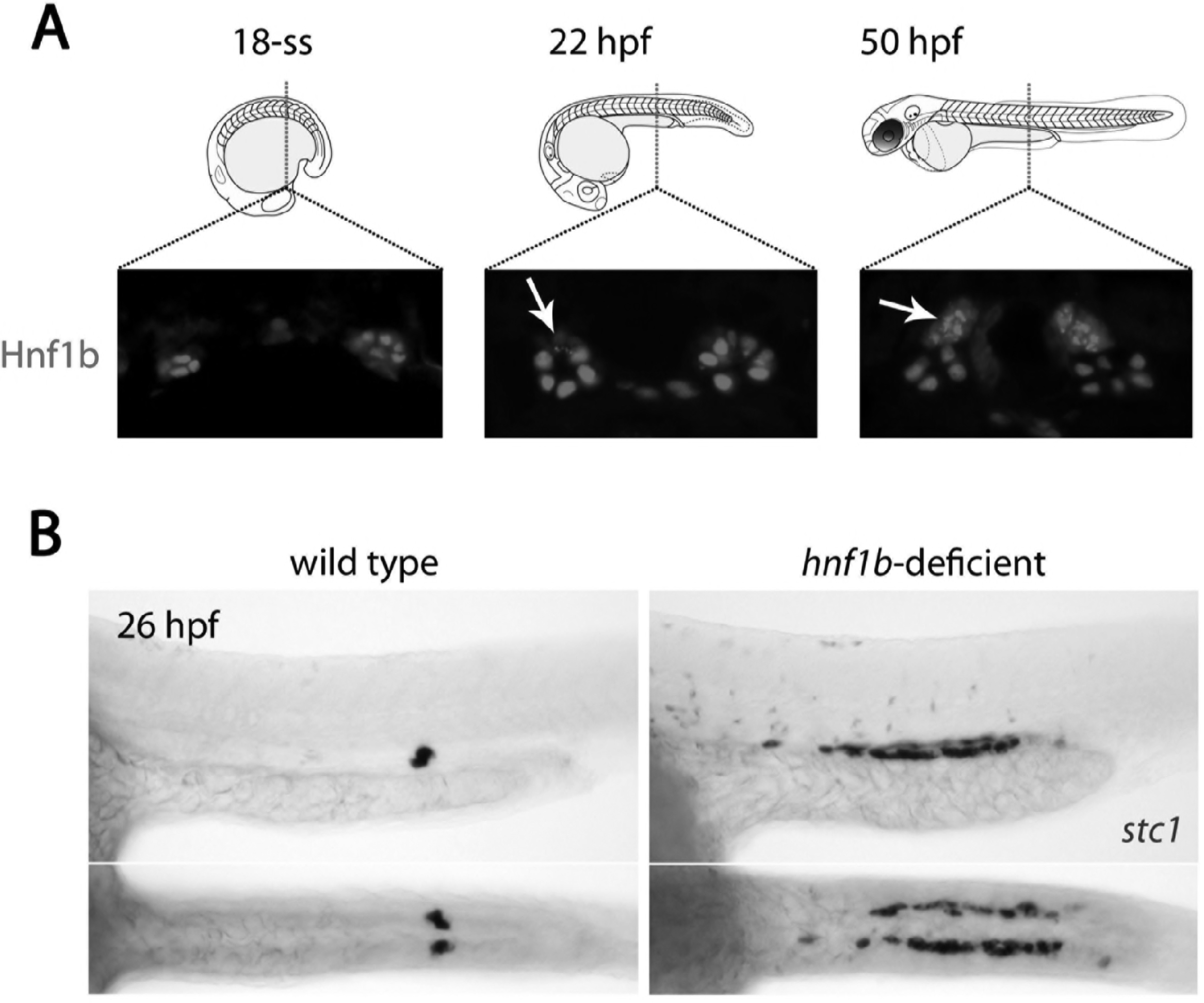
Hnf1b is translocated to the cytoplasm in prospective CS cells. A) Panels show Hnf1b immunostains (red) on transverse cryosections of zebrafish embryos through the region of the trunk where CS glands form at 22 hpf, 36 hpf and 50 hpf. Arrows indicate CS cells that have undergone nuclear export of Hnf1b. B) Top panels show lateral views and bottom panels show dorsal views of wild-type and *hnf1b*-deficient embryos stained for *stc1*.

### Irx3b inhibits CS cell fate commitment in the pronephros

The formation of the DE segment has previously been shown to be regulated by the Irx3b transcription factor, which acts downstream of Hnf1b (R. A. Wingert & Davidson, 2011). We therefore explored the involvement of Irx3b in CS gland formation in loss-of-function studies. We found that *irx3b* knockdown using morpholinos did not prevent the initial formation of the DE segment, as *slc12a1* displays a normal expression pattern in *irx3b* morphants at 18 hpf (n=21) and 20 hpf (n=24, Fig 4A). However, we noted that the length of the DE segment was reduced in rx3b-deficient embryos at 22 hpf (n=23) and 24 hpf (n=25, Fig.4A) and there was a corresponding expansion in *stc1*^+^ CS cells from 24 hpf onwards (Fig.4B). By 36 hpf, the DE region lacked transcripts for *slc12a1*, consistent with previous reports (R. A. Wingert & Davidson, 2011) and *cdh17* was down-regulated in the region where ectopic *stc1*^+^ cells are present (Fig.4B). An analysis of live *Tg(cdh17:egfp)* embryos that are deficient in *irx3b* showed that these ectopic CS cells still underwent apical constriction but were not fully extruded from the tubule (Fig.5A). In addition, we found that the tubule cells rostral to the enlarged CS gland display a stretched morphology in *irx3b*-deficient animals, both in live *Tg(cdh17:egfp)* embryos (Fig.5A) and by expression analysis of the pan-tubule marker *atp1a1a.4* (Fig.5B). Conversely, the DL segment was compacted and shorter in *irx3b*-depleted animals (Fig.5A and 5B).

**Figure 4:**
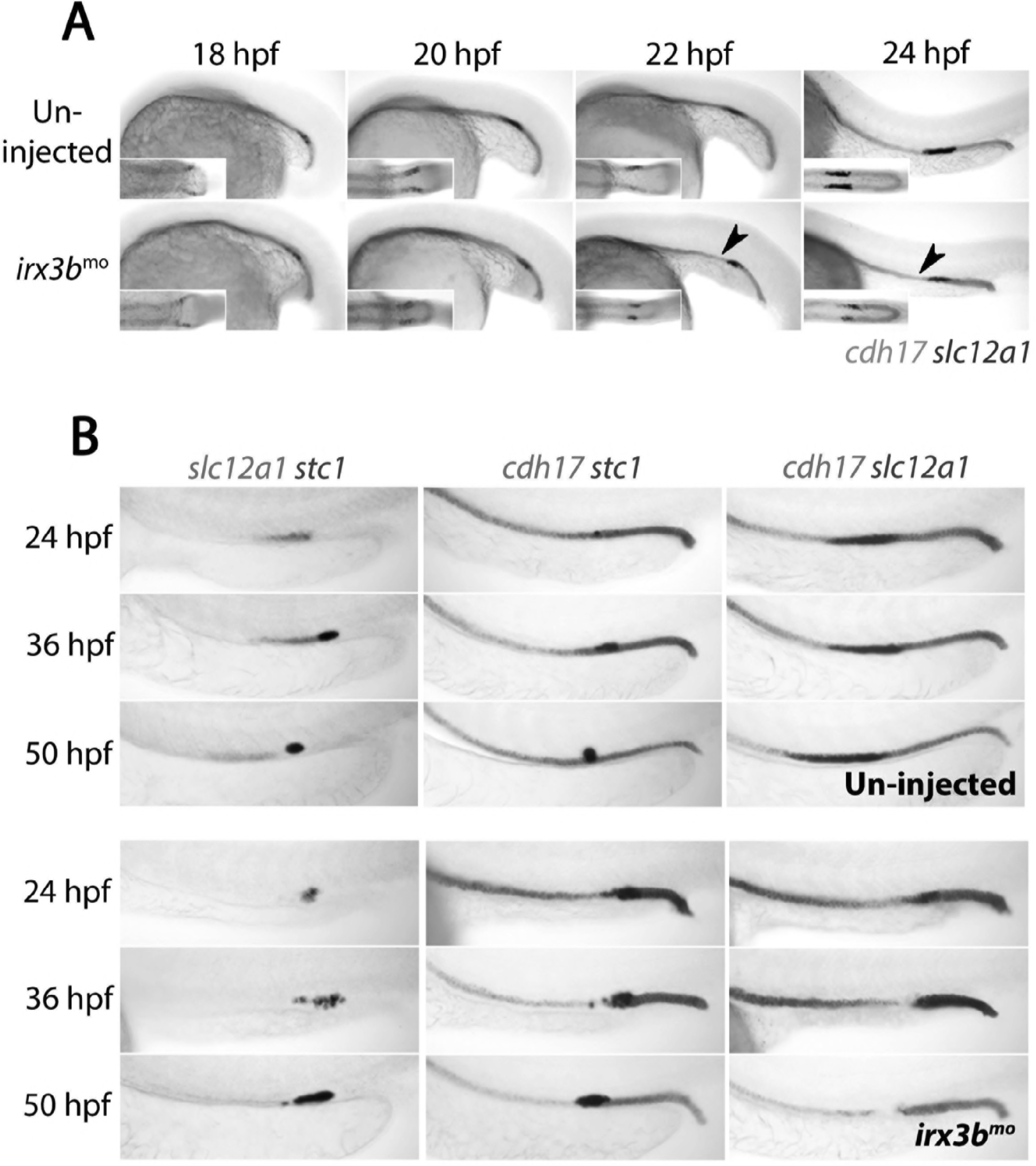
*irx3b* depletion causes the entire DE segment to adopt a CS fate. A) Panels show lateral views of 18 hpf, 20 hpf, 22 hpf and 24 hpf wild-type and *irx3b* morphant embryos co-stained for *cdh17* (red) and *slc12a1* (purple). Black arrowheads highlight the initiation of pronephric phenotypes in *irx3b* morphants. B) Panels show lateral views of control un-injected embryos and *irx3b* morphants double stained for either *slc12a1(red)*/ *stc1*(purple), *cdh17*(red)/ *stc1*(purple) or *cdh17*(red)/ *slc12a1*(purple) at 24 hpf, 36 hpf and 50 hpf.

**Figure 5:**
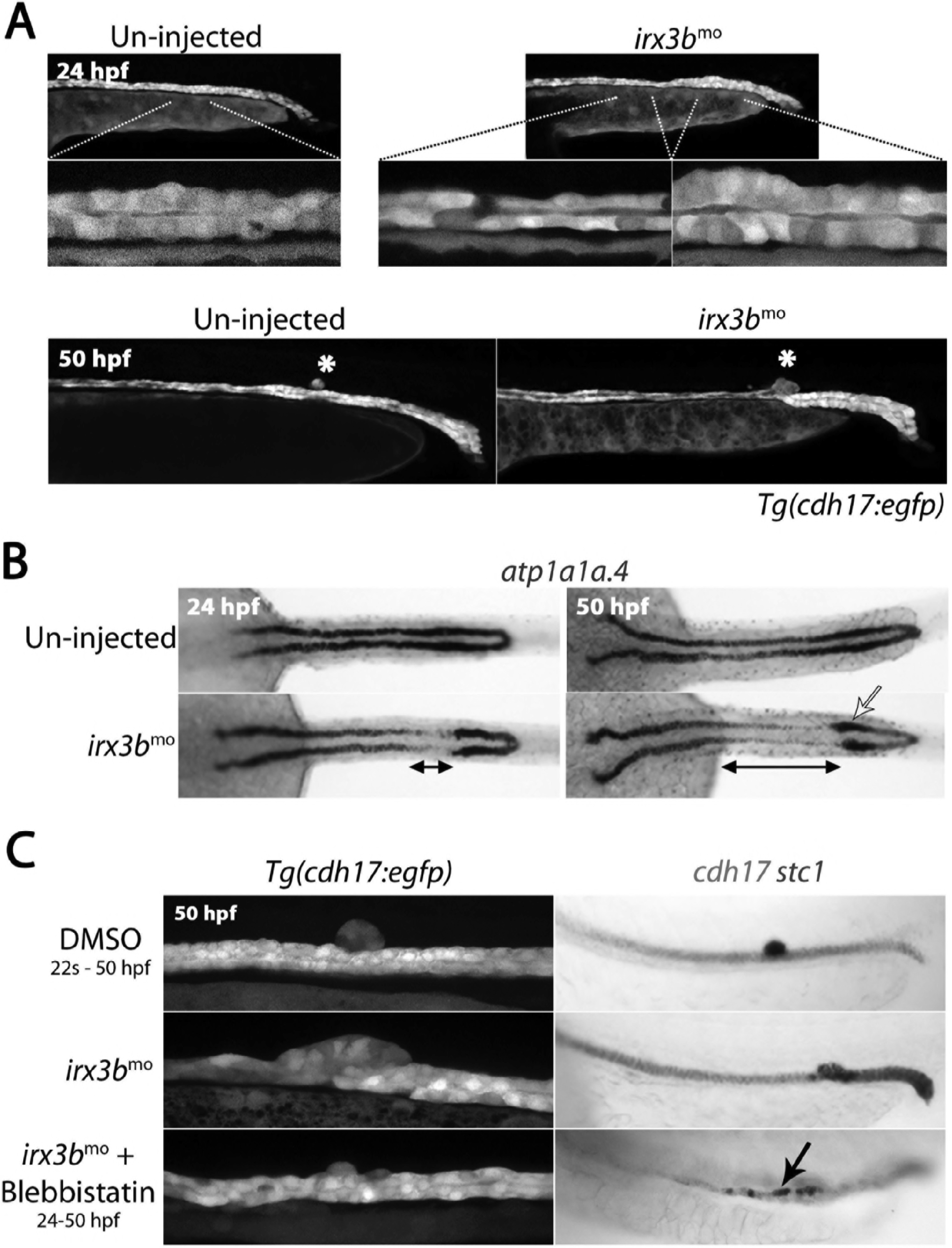
*irx3b* depletion causes aberrant pronephric morphogenesis. A) Lateral views of live un-injected and *irx3b* morphant *Tg(cdh17::egfp)* embryos with lower panels showing higher magnification of indicated pronephric regions. B) Panels show dorsal view images of embryos stained for *atp1a1a.4* at the stages indicated. Double-ended black arrows highlight areas of reduced *atp1a1a.4* expression immediately anterior to a thickened distal tubule (yellow arrow) at 24 hpf and 50 hpf in *irx3b* morphants. C) Lateral views of the trunk of *Tg(cdh17:egfp)* embryos (left panels) and embryos double stained for *cdh17* (red) and *stc1* (purple; right panels) after the indicated treatments. Black arrow highlights the ectopic *stc1*^+^ cells that remain in the tubule after *irx3b* knock down and Blebbistatin treatment.

Given that the tubule starts moving rostrally by 29 hpf (Vasilyev et al., 2009), the morphological effects caused by *irx3b* knockdown suggests that an enlarged and un-extruded CS gland may prevent the DL segment from undergoing this movement, resulting in greater stretching of the more proximal tubule cells and compaction of the distal tubule. To test this, *irx3b*-deficient animals were treated with Blebbistatin to block apical constriction. These animals were found to display relatively normal tubule morphology with ectopic *stc1*^+^ cells scattered throughout the tubule (80%, n=25, Fig.5C). Taken together, these results suggest that Irx3b inhibits the transdifferentiation of DE cells to a CS fate and that the failed extrusion of the ectopic CS cells seen in *irx3b*-deficient embryos alters tubular cell morphology due to a block in the rostral migration of the DL segment.

### Irx3b suppresses CS cell formation

We then went on to examine a link between Irx3b and Notch, as the Irx factors have been implicated as both positive and negative regulators of Notch signalling (Dominguez & de Celis, 1998; Scarlett, Pattabiraman, Barnett, Liu, & Anderson, 2015). In addition, Notch has recently been linked to CS gland development (B. E. Drummond, Li, Marra, Cheng, & Wingert, 2016) and Notch components are expressed in CS cells (Fig.S4). We first analysed the *mindbomb* mutant, which is defective in Notch signalling (Itoh et al., 2003; Koo et al., 2005; Schier et al., 1996), and found that fewer *stc1*^+^ cells form in mutants (Fig.6A). Similarly, treatment of zebrafish embryos with the γ-secretase inhibitor compound E (cpdE) from 18 hpf onwards also reduced the number of *stc1*^+^ cells (Fig.6B). We next performed epistasis experiments to uncover if Notch signals are required upstream or downstream of *irx3b*. In our hands, we found that the effective dose of the *irx3b* morpholino was slightly toxic to the embryo, suggesting it may have off-target effects. These effects were exaggerated in embryos treated with γ-secretase inhibitors (data not shown). To minimise these effects and to also confirm the specificity of the *irx3b* morpholino knockdown phenotype, we utilised the CRISPR-Cas9 system to introduce mutations into the *irx3b* gene (Burger et al., 2016; Gagnon et al., 2014). Injection of two gRNAs targeting *irx3b* reproduced the *irx3b* morpholino phenotype in G_0_ animals (crispants) by increasing the number of *stc1*^+^ cells in the DE region of the tubule (n=45/49, Fig.6B). T7 endonuclease 1 analysis (Huang, Cheong, Lim, & Li, 2012) for DNA mismatches in *irx3b* crispants indicated successful mutagenesis was achieved (Fig.S5). These crispant embryos were also grown to adulthood and used to generate stable mutant lines that similarly phenocopied the morphant/ crispant phenotypes (Fig.S6). In *irx3b* crispants treated with cpdE, the number of *stc1*^+^ cells was robustly reduced (n=37/37). In conclusion, these results suggest that Irx3b acts upstream or in parallel to Notch signalling to inhibit the transdifferentiation of DE cells into CS cells.

**Figure 6:**
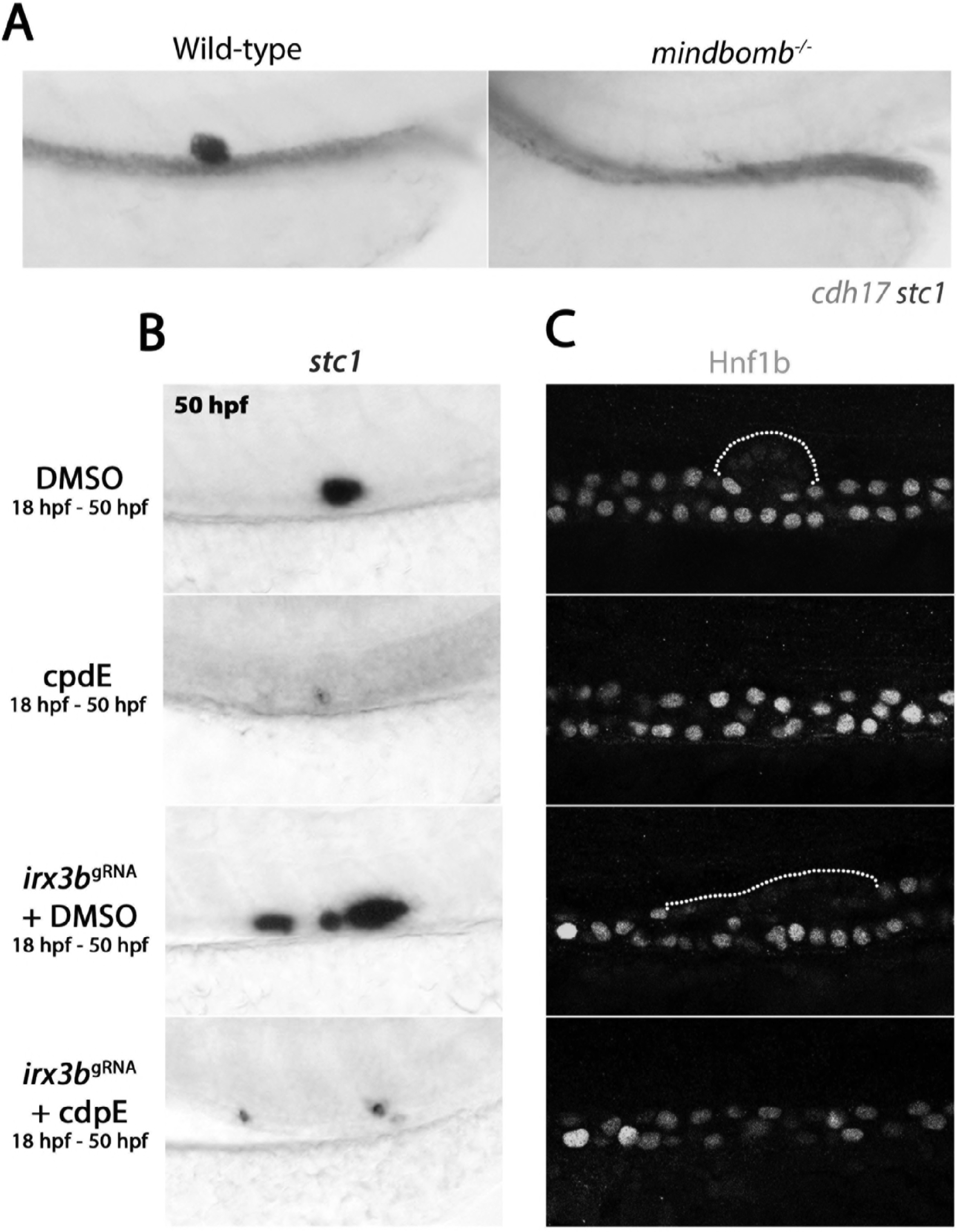
Notch inhibition precludes CS gland formation. A) Panels show lateral views of control un-injected embryos and *mindbomb* mutants double stained for *cdh17*(red)/ *stc1*(purple) 50 hpf. B) Lateral views of the posterior trunk showing *stc1* expression after the indicated treatments. C) Lateral views of the posterior trunk showing Hnf1b staining after the indicated treatments.

We next analysed the effects of *irx3b* depletion and γ-secretase inhibition on the intracellular localisation of Hnf1b. We found in *irx3b* crispants that the number of cells with nuclear-localised Hnf1b was qualitatively decreased in the region of the DE segment (Fig.6C). In contrast, in wild-type (n=27/27) or *irx3b* crispant embryos that had been treated with cpdE n=25/25), Hnf1b was retained in the nucleus of renal tubule cells in the DE region (Fig.6C). These results are consistent with a model in which Notch signalling and the cytoplasmic sequestration of Hnf1b in a subset of DE epithelial cells leads to the transdifferentiation of renal epithelial cells into CS cells.

## Discussion

In this study we have discovered a rare example of naturally occurring transdifferentiation that occurs in the zebrafish embryonic kidney and results in the formation of endocrine cells from the renal tubular epithelium. By live-cell imaging we show that the CS cells are extruded from the tubule via apical constriction and mass cell extrusion. We investigated the molecular pathways governing this transdifferentiation event and identified the Notch pathway as being a critically important signal together with cytoplasmic sequestration of Hnf1b, a master determinant of renal epithelial fate. The Irx3b transcription factor was found to play an inhibitory role in this process by preventing widespread transdifferentiation of the DE segment. These results establish a new model system to understand the molecular and cellular processes governing cellular plasticity *in vivo*.

Our data indicate that the transdifferentiation of DE to CS fate occurs in sequential steps and we provide both genetic (*mindbomb* mutants) and chemical (CpdE) evidence to implicate Notch signalling as an early critical event. Unexpectedly, these findings contradict another report in which global overexpression of *nicd1*, encoding the constitutively active signaling moiety of the Notch 1 receptor, decreases the number of CS cells and treatment with the DAPT Notch inhibitor was found to increase CS cell number (B. E. Drummond et al., 2016). In this prior study, Notch was activated at 16 hpf while it was inhibited from late gastrulation onwards using DAPT. In our study we used CpdE and treated from 18 hpf. Thus, the discrepancy in outcomes may be related to the efficacy of the inhibitors used as well as the onset/ duration of when the pathway is modulated, particularly given that Notch signaling is known to be dose responsive and to serve multiple functions at different times during development (Guruharsha, Kankel, & Artavanis-Tsakonas, 2012).

Notch has well-described roles in determining binary cell fate decisions during progenitor cell decisions including in the zebrafish pronephros where it acts by a classical lateral-inhibition mechanism to inhibit a multi-ciliated phenotype in renal progenitors (Liu, Pathak, Kramer-Zucker, & Drummond, 2007; Ma & Jiang, 2007; Marra & Wingert, 2016). A similar lateral-inhibition pathway may be required for the DE-to-CS transdifferentiation but with the timing of this occurring after the multi-ciliated cell fate decision (~16 hpf) (Liu et al., 2007; Ma & Jiang, 2007; Rebecca A Wingert et al., 2007). We, and others, have found that a number of Notch signalling components are expressed in the pronephric tubules during the period that the CS gland arises (Liu et al., 2007; Ma & Jiang, 2007; Rebecca A Wingert et al., 2007). A comprehensive loss-of-function analysis of these genes is needed in order to determine which of these is involved in CS formation. In the context of transdifferentiation, work in *C. elegans* has implicated Notch in the conversion of rectal epithelial Y cells into motor neurons, although in this case the Y cell transitions to an intermediary cell state rather than directly transdifferentiating as seen in our study (Jarriault et al., 2008). In a mammalian example, bronchial cells known as ‘club’ cells, which depend on active Notch for their maintenance, switch to a ciliated phenotype following disruption of Notch signalling in adult mice, without an apparent dedifferentiation stage or need for cell division (Lafkas et al., 2015). In this example, ‘club’ cells and ciliated cells share a common progenitor during lung development whereas in our study the renal and glandular lineages have, until now, been considered to be developmentally distinct fates. Indeed, the conversion we see of a renal epithelial cell becoming an endocine gland can be considered more of a trans-organogenesis event.

Mechanistically, how Notch signalling induces the DE to CS fate switch requires more study. Our observation that the Hnf1b transcription factor, a master regulator of renal epithelial fate, becomes sequestered in the cytoplasm of cells undergoing transdifferentiation suggests that Notch regulates the levels of Hnf1b in the nucleus either by inducing Hnf1b nuclear export or alternatively, if the nuclear turnover of Hnf1b is high, by preventing newly translated Hnf1b from entering the nucleus. We favour the former notion as Hnf1b is considered a relatively stable ‘bookmarking’ factor that remains associated with chromatin during cell division where it reinstates transcriptional programs following mitotic exit (Verdeguer et al., 2010). As Hnf1b is critical for renal tubule differentiation in zebrafish and mice (Heliot et al., 2013; Massa et al., 2013; Naylor & Davidson, 2014; Naylor et al., 2013), we hypothesize that there is an on-going requirement for Hnf1b in the tubule to maintain renal epithelial identity and that sequestration of this factor is sufficient to induce the transdifferentiation of DE cells into CS cells. In support of this, we found that *hnf1b*-deficient embryos show ectopic *stc1* expression along a broad stretch of the pronephric tubule, consistent with a CS program being activated in these cells. Whether CS identity is a default state in the tubule is unclear, but we have observed that the Notch ligand *jagged-2a* is abnormally maintained in the pronephric tubule of *hnf1b*-deficient embryos (data not shown) consistent with aberrant Notch signalling also being involved in the induction of ectopic *stc1*. The notion that sequestration of a master transcription factor can induce transdifferentiation provides a new paradigm for the field, as most studies of cell fate plasticity have reported a need for regulatory factors to be up-regulated in order to drive the process, particularly those involving non-physiological ‘reprogramming’ (reviewed in Daniel, Lemischka, & Moore, 2016).

Our data point to the Irx3b transcription factor as playing an important role in the DE-to-CS fate switch. We have previously linked Hnf1b and Irx3b together by demonstrating that *hnflb*-deficient embryos fail to express *irx3b* in the renal tubules (Naylor et al., 2013), placing *hnf1b* upstream of *irx3b* in a genetic hierarchy. In this study we show that *irx3b* deficiency does not influence the initial specification of the DE segment as previously thought (R. A. Wingert & Davidson, 2011) but instead leads to the dramatic transdifferentiation of this segment into CS cells. Consistent with our notion that Hnf1b sequestration drives this transdifferentiation, we found that the ectopic CS cells display cytoplasmic Hnf1b. Furthermore, inhibiting Notch signalling rescued the CS expansion phenotype in *irx3b*-deficient animals, including Hnf1b nuclear localisation, placing Irx3b upstream or in parallel to the Notch pathway. Irx/ Iroquois family transcription factors have been reported to modulate Notch signalling in other cellular contexts. Most notably during *Drosophila* eye development, where the Iroquois genes establish where Notch activation occurs by controlling the levels of Fringe, a glycosyltransferase that alters the sensitivity of the Notch receptors to transactivation from the different classes of Notch ligand (Moloney et al., 2000; Panin, Papayannopoulos, Wilson, & Irvine, 1997). In vertebrates, including zebrafish, there are three Fringe homologues (Qiu, Xu, Haddon, Lewis, & Jiang, 2004) but their expression patterns during CS formation have not been closely examined. One speculative idea is that Irx3b modulates Notch signalling in the DE segment via the regulation of *fringe* or other Notch pathway genes, to ensure that the CS-inducing effects of Notch signalling are suppressed. This may occur by promoting one Notch-Ligand pairing over another or by altering the strength of the Notch signal. It is also interesting to note that cross-regulatory interactions between Notch and Irx3 have been shown in human microvascular endothelial cells where *Irx3* induces expression of *Delta-like ligand 4* but is in turn repressed by the Notch mediator Hey1 via direct binding to the *Irx3* promoter (Scarlett et al., 2015). How the proposed repressive effects of Irx3b on Notch signalling are overcome so that the DE-to-CS transdifferentiation event can be initiated at the appropriate time during development remain to be determined in future studies. Similarly, how the Notch, Irx3b and Hnf1b factors interact with other transcriptional regulators that have been reported to control CS cell number, such as Sim1a and Tbx2a/b (Cheng & Wingert, 2015; B. E. Drummond et al., 2016), also need investigating.

The second key step in the transdifferentiation of DE cells to a CS gland is the expulsion of the fate-changed cells from the basal side of the tubule. This process presents a challenge to the embryo as any loss of cells from the epithelium has the potential to compromise the barrier function of the tubule and cause lumenal fluid to leak out into the interstitium. Our live-cell imaging revealed that the first step in achieving this feat is the constriction of the apical membranes of the CS cells. Apical constriction is a well-known feature of polarised epithelia that drives bending, folding and tube formation during the formation of different tissues and organs (Martin & Goldstein, 2014; A. C. Martin, M. Kaschube, & E. F. Wieschaus, 2009; Mason, Tworoger, & Martin, 2013; Sadler, 2005; Sawyer et al., 2010). Apical constriction is driven by contraction of an apical meshwork of F-actin by the molecular motor myosin IIA (Adam C. Martin, Matthias Kaschube, & Eric F. Wieschaus, 2009) and consistent with this, we found that blebbistatin treatment inhibited CS cell extrusion. Interestingly, in the *Drosophila* eye-antennal disc, Notch activation has been found to trigger actomyosin-mediated cell apical constriction, raising the possibility that the Notch pathway may both induce DE-to-CS transdifferentiation and initiate apical constriction (Ku & Sun, 2017).

In the developing avian lung, where the tubular epithelium of the primary bronchus undergoes apical constriction in localised areas, the result is the formation of lung buds that then grow out as new airways (Locy WA, 1916). Unlike this system, the ‘bud’ of CS cells we observe is made up of far fewer cells, does not transition to a tubular outgrowth and proliferation is not required for its extrusion, although we do observe CS cells proliferating once they are free from the tubule. Instead, the budding CS cells are ‘shed’ basally, *en masse*, as an intact ball of cells. Thus, we conclude that while apical constriction initiates the process, additional cellular behaviours must then be required to extrude the cells. One possible mechanism for CS cell extrusion is that the CS cells undergo EMT in order to disassemble their tight junctions with their neighbours. This is an attractive hypothesis, particularly given that this process operates in other related tissues such as the pancreas, where the primitive duct epithelium responds to low Notch signalling to down-regulate Cdh1 in single cells that then undergo EMT and delaminate as endocrine precursors (Shih et al., 2012). However, we do not observe a loss of Cdh1, which is a widely used definition of EMT (Kalluri & Weinberg, 2009). Furthermore, from our histological analysis we have not been able to capture an intermediary mesenchyme-like state and observe no migration of individual CS cells. Instead, we find that CS cells rapidly and seamlessly change from a cuboidal epithelium in the renal tubule to a simple lobular epithelium of the CS gland. Based on this we favour a model in which CS cells are extruded by a non-EMT-based pathway that may be more comparable to the epithelial remodelling that occurs during primary spinal cord formation in the embryo. Here, the neural plate undergoes apical constriction and invaginates before pinching off as the neural tube while the surrounding non-neural ectoderm fuses together (Nikolopoulou, Galea, Rolo, Greene, & Copp, 2017). The exact molecular mechanism for how barrier function is maintained during neural tube closure is not fully understood but epithelial-to-mesenchymal (EMT) transitions appear suppressed (Ray & Niswander, 2016).

Prior work has shown that overcrowding of epithelial sheets results in live cell extrusion of single cells by a mechanism that involves the formation of an actomyosin contractile ring in cells surrounding the extruded cell. This ring then contracts and the innermost cell is pushed out of the epithelium (Eisenhoffer et al., 2012; Gu et al., 2011). We performed laser ablations immediately anterior and posterior to the CS gland to disrupt such a ring in neighbouring cells but this failed to prevent CS extrusion. In addition, we have found that treating embryos with the S1P_2_ receptor inhibitor JTE-013 has no effect on CS extrusion (data not shown). Thus, it is unlikely that this mechanism of live cell extrusion is playing a role in CS gland shedding.

Taken together, our data show that the conversion of DE cells to a CS gland fate in the zebrafish renal tubule represents a novel example of a developmentally programmed direct transdifferentiation event coupled to an unusual live cell extrusion behaviour. This model provides new ways to interrogate the plasticity of differentiated cellular states, will advance efforts to harness transdifferentiation for therapeutic applications, and furthers our understanding of epithelial cell behaviours.

## Methods and Materials

### Zebrafish husbandry

Zebrafish embryos were maintained and staged according to established protocols (Kimmel, Ballard, Kimmel, Ullmann, & Schilling, 1995) and in accordance with the University of Auckland’s Animal Ethics Committee (protocol 001343). Paired matings of wild-type *Tübingen (Tg)*, transgenic *Tg(cdh17:egfp)*, mutant *mindbomb*^*+/−*^ and mutant *Tübingen*^*+/hi1843*^ adults were carried out in order to collect embryos used to perform the experiments described.

### MO design and injection

Previously validated morpholinos to *irx3b* (5′- ACCGGGAGGACTGCGGGGAAACTCG -3′ (R. A. Wingert & Davidson, 2011)) and *hnf1bb* (5’-CTTGGACACCATGTCAGTAAA-3’ (Choe, Hirsch, Zhang, & Sagerstrom, 2008)) were purchased from Gene Tools LLC and resuspended in 1X Danieau solution. One-cell embryos were injected with 1-5 nl of morpholino at 1.5 ng/nl for *irx3b* and 5 ng/nl for *hnf1bb*. Embryos were incubated at 28.5°C to the desired stage and fixed in 4% paraformaldehyde. All experiments were repeated at least 3 times to reproducible yield the phenotypes as described in the text.

### gRNA design and injection

gRNAs were designed using CHOPCHOP (Harvard) software and synthesized according to published protocols (Gagnon et al., 2014). We modified this protocol slightly to create two gRNAs that simultaneously targeted the first exon (5’-GGCGCGGAGATCTCGGTCAC-3’) and second exon (5’-GGATGCAGAAAAACGAGATG-3’) of the *irx3b* locus in order to increase the chance of biallelic gene disruption. SP6 promoter containing gRNA templates were generated using Phusion polymerase (ThermoFisher). Transcripts were synthesised using the SP6 Megascript kit (Ambion). Cas9 protein that included a nuclear localisation signal was purchased from PNA-BIO Inc. (CP02). Approximately 10 nl of 0.2 ng/μl *irx3b* gRNAs^1+2^, 0.4 ng/μl Cas9 and 300 mM KCl were injected directly into the cell of early one-cell stage embryos as per the Burger, A. *et al* (2016) protocol. For the T7 endonuclease assay, 5 uninjected and 5 *irx3b* gRNAs injected embryos were lysed in 50 μl of 50 mM NaOH at 95°C for 20 minutes. 5 μl of 100 mM Tris HCl pH 8.5 was added to the lysed embryos and the solution was homogenised. 1 μl of the lysed embryos containing genomic DNA was used in a PCR reaction to amplify *irx3b* exon 1, with PCR primers flanking the site of mutagenesis (Forward primer 5’-TCCCGCAGCTAGGCTATAAGTA-3’; Reverse primer 5’-ACGGGATCAAATCTGAGCTATT-3’). After purification of the PCR product, 200 ng of amplified DNA, 2 μl of 10X NEB buffer and water (to a final volume of 19 μl) was mixed together and placed in a thermocycler for rehybridisation; 5 minutes at 95°C, ramp down to 85°C at −2°C/second, ramp down to 25 °C at −0.1°C/second, 25 °C for 10 minutes. 1 μl of T7 Endonuclease 1 (NEB) was then added to the rehybridisaed PCR product and incubated at 37°C for 20 minutes before the products were ran on a 1.5% agarose gel to analyse for digestion.

### Drug treatments

Embryos were dechorionated and exposed to pharmacological agents at the time-points outlined in the text. Blebbistatin (SigmaAldrich #B0560) and Compound E (Abcam, ab142164) were dissolved in DMSO to a stock concentration of 10 mM and diluted in E3 media to a working concentration of 10 μM and 50 μM, respectively. Aphidicolin (SigmaAldrich #A0781) was dissolved to a 10 mM stock solution with DMSO. The combined Hydroxyurea (20 mM) and Aphidicolin (150 μM) solution (termed HUA in the text) was made up in E3 embryo medium.

### Whole mount in situ hybridisation and immunohistochemistry

Whole mount *in situ* hybridization was performed using protocols previously described (Thisse & Thisse, 2008). Digoxigenin and Fluorescein anti-sense riboprobes were synthesized using T7/T3/SP6 RNA polymerase transcription kits (Roche Diagnostics) from plasmids used previously (R. A. Wingert & Davidson, 2011; Rebecca A Wingert et al., 2007). For double *in situ* hybridizations, alternative Dig- or Flu- riboprobes were used and alkaline phosphatase was inactivated by two 15 minute 100 mM Glycine (pH 2.2) treatments. To perform whole mount antibody staining, fixed embryos were washed twice in PBS containing 0.05% Tween20 (PBST), then placed in PBS containing 0.5% Triton X100 for 20 minutes to permeabilize the embryo. Embryos were then washed twice in TBS containing 0.05% Triton X100 (TBST) and blocked in TBST containing 3% BSA and 5% Goat serum for at least one hour. The Hnf1b (SantaCruz Ab #22840) and E-cadherin (SantaCruz Ab #7870) primary antibodies was added at 1:500 dilution overnight at 4°C. Embryos were washed 5 times in PBST before Goat anti-Rabbit IgG DyLight594 conjugated secondary antibody (Abcam #96901) was added at 1:500 dilution. Embryos were incubated in the secondary antibody for 2 hours at room temperature, washed twice in PBST then imaged on a FV1000 LiveCell confocal microscope (Olympus).

### Cryosectioning

Zebrafish embryos were fixed in 4% PFA overnight at 4°C and washed twice in PBS and then placed in 1% low melting point agarose containing 5% sucrose and 0.9% agar (made up in water) in cryomoulds. After one hour the cryoblocks were removed from the cryomoulds and placed in 30% Sucrose/PBS. Cryoblocks were incubated overnight at 4°C then 30% Sucrose/PBS was replaced and left for another night at 4°C. Cryoblocks were flash frozen on dry ice and immediately sectioned. 14 μm sections were cut in a cryostat machine set to −25°C and transferred to gelatin-coated microscope slides.

### EdU assay

For exposure to the EdU label, *Tg(cdh17:egfp)* embryos were injected with 0.5 μg/ml EdU (SantaCruz #284628). Injections were performed into the yolk (embryo was immobilized in a well formed from a 1% agarose bed in a 45 mm petri dish) and left in E3 embryo medium for the periods of development outlined in the text (we find incubating embryos in E3 media containing EdU only permitted superficial incorporation at post gastrula stages). Embryos were fixed at the desired stage and then incorporated EdU was detected using a Click-iT EdU Alexa Fluor 594 kit (Life Technologies). We found that this labelling quenched the endogenous eGFP fluorescence in the transgenic embryo, thus antibody staining using the same protocol described above was performed for eGFP (SantaCruz Ab #8334, 1:500).

### Time-lapse imaging

32 hpf *Tg(cdh17:egfp)* embryos were anaesthetised in embryo media containing 0.02% Ethyl 3-aminobenzoate methanesulfonate (Tricaine) and 0.003% N-phenylthiourea (PTU). Anaesthetised embryos were placed in a well made in a 1.5% agarose gel and positioned appropriately for image capture. Z-stack confocal images were captured every 20 minutes for 20 hours on a FV1000 LiveCell confocal microscope (Olympus) to generate the time-lapse video.

## Acknowledgements

We thank S. Patke for managing and taking care of the fish. This work was supported by a Health Research Grant of New Zealand 13/332 Alan J. Davidson.

---

**Supplementary Figure 1: Cell division is not required for CS formation** A) Panels show lateral views of *Tg(cdh17:egfp)* embryos stained for EdU labelling at 36 hpf (left panel) and 50 hpf (two right panels) after treatment with EdU between the stages shown. B) Lateral views of wild-type and HUA treated animals at 50 hpf. Left panels are lateral views of *Tg(cdh17:egfp)* and right panels are lateral views of embryos stained for *cdh17* (red) and *stc1* (purple) expression.

**Supplementary Figure 2: *hnf1ba* is not expressed in the CS gland** Panel shows a 50 hpf wild-type embryo stained for *hnf1ba*. Black arrow indicated the approximate position of the CS gland.

**Supplementary Figure 3: Hnf1b nuclear export at 50 hpf** Lateral view of a 50 hpf *Tg(cdh17:egfp)* (green) embryo immunostained for Hnf1b (red) and counterstained for DAPI (blue). Hnf1b in the CS was non-nuclear (white arrow), in contrast to nuclear Hnf1b in the tubules (black arrow).

**Supplementary Figure 4: Notch components are expressed in the CS gland** Lateral views of wild-type embryos stained for *notch3* and *jagged2b* (*jag2b*) at the stages shown, black arrows indicate expression of notch factors in the CS gland.

**Supplementary Figure 5: T7 endonuclease assay on *irx3b* crispants shows efficient mutagenesis** Panel shows gel image of *irx3b* PCR product amplified from genomic DNA and processed through the T7 endonuclease 1 protocol (see methods section).

**Supplementary Figure 6: *irx3b* crispants and stable mutants recapitulate the renal phenotypes associated with *irx3b* morpholino knock down** Panels show lateral views of *Tg(cdh17:egfp)* embryos in wild-type, *irx3b* morphant, *irx3b* crispant and germline stable *irx3b* mutant embryos.

**Supplementary Movie 1: Time-lapse imaging showing extrusion of the CS endocrine gland from the pronephric tubule** Movie shows a lateral view of the posterior trunk region of a Tg(cdh17:egfp) embryo between 32 hpf and 48 hpf. Green fluorescent cells label the pronephric tubule and the CS cells, with the latter observed as a bulge of cells that undergo extrusion during the time-lapse.

## References

Blaine, J., Chonchol, M., & Levi, M. (2015). Renal control of calcium, phosphate, and magnesium homeostasis. Clin J Am Soc Nephrol, 10(7), 1257–1272. doi:10.2215/cjn.09750913

Burger, A., Lindsay, H., Felker, A., Hess, C., Anders, C., Chiavacci, E., … Mosimann, C. (2016). Maximizing mutagenesis with solubilized CRISPR-Cas9 ribonucleoprotein complexes. Development, 143(11), 2025–2037. doi:10.1242/dev.134809

Cheng, C. N., & Wingert, R. A. (2015). Nephron proximal tubule patterning and corpuscles of Stannius formation are regulated by the sim1a transcription factor and retinoic acid in zebrafish. Dev Biol, 399(1), 100–116. doi:10.1016/j.ydbio.2014.12.020

Choe, S. K., Hirsch, N., Zhang, X., & Sagerstrom, C. G. (2008). hnf1b genes in zebrafish hindbrain development. Zebrafish, 5(3),179–187. doi:10.1089/zeb.2008.0534

Cohen, R. S., Pang, P. K., & Clark, N. B. (1975). Ultrastructrue of the Stannius corpuscles of the killifish, Fundulus heteroclitus, and its relation to calcium regulation. Gen Comp Endocrinol, 27(4), 413–423.

Daniel, M. G., Lemischka, I. R., & Moore, K. (2016). Converting cell fates: generating hematopoietic stem cells de novo via transcription factor reprogramming. Annals of the New York Academy of Sciences, 1370(1), 24-35. doi:10.1111/nyas.12989

Dominguez, M., & de Celis, J. F. (1998). A dorsal/ventral boundary established by Notch controls growth and polarity in the Drosophila eye. Nature, 396(6708), 276–278. doi:10.1038/24402

Drummond, B. E., Li, Y., Marra, A. N., Cheng, C. N., & Wingert, R. A. (2016). The tbx2a/b transcription factors direct pronephros segmentation and corpuscle of Stannius formation in zebrafish. Dev Biol. doi:10.1016/j.ydbio.2016.10.019

Drummond, I. A., & Davidson, A. J. (2010). Zebrafish kidney development. Methods Cell Biol, 100, 233–260. doi:10.1016/b978-0-12-384892-5.00009-8

Drummond, I. A., Majumdar, A., Hentschel, H., Elger, M., Solnica-Krezel, L., Schier, A. F., … Fishman, M. C. (1998). Early development of the zebrafish pronephros and analysis of mutations affecting pronephric function. Development, 125(23), 4655–4667.

Eisenhoffer, G. T., Loftus, P. D., Yoshigi, M., Otsuna, H., Chien, C. B., Morcos, P. A., & Rosenblatt, J. (2012).Crowding induces live cell extrusion to maintain homeostatic cell numbers in epithelia. Nature, 484(7395), 546–549. doi:10.1038/nature10999

Fang, D., Nguyen, T. K., Leishear, K., Finko, R., Kulp, A. N., Hotz, S., … Herlyn, M. (2005). A tumorigenic subpopulation with stem cell properties in melanomas. Cancer Res, 65(20), 9328–9337. doi:10.1158/0008-5472.can-05-1343

Gagnon, J. A., Valen, E., Thyme, S. B., Huang, P., Akhmetova, L., Pauli, A., … Schier, A. F. (2014). Efficient mutagenesis by Cas9 protein-mediated oligonucleotide insertion and large-scale assessment of single-guide RNAs. PLoS One, 9(5), e98186. doi:10.1371/journal.pone.0098186

Gu, Y., Forostyan, T., Sabbadini, R., & Rosenblatt, J. (2011). Epithelial cell extrusion requires the sphingosine-1-phosphate receptor 2 pathway. J Cell Biol, 193(4), 667–676. doi:10.1083/jcb.201010075

Guruharsha, K. G., Kankel, M. W., & Artavanis-Tsakonas, S. (2012). The Notch signalling system: recent insights into the complexity of a conserved pathway. Nat Rev Genet, 13(9), 654–666. doi:10.1038/nrg3272

Heliot, C., Desgrange, A., Buisson, I., Prunskaite-Hyyrylainen, R., Shan, J., Vainio, S., … Cereghini, S. (2013). HNF1B controls proximal-intermediate nephron segment identity in vertebrates by regulating Notch signalling components and Irx1/2. Development, 140(4), 873–885. doi:10.1242/dev.086538

Horsfield, J., Ramachandran, A., Reuter, K., LaVallie, E., Collins-Racie, L., Crosier, K., & Crosier, P. (2002). Cadherin-17 is required to maintain pronephric duct integrity during zebrafish development. Mech Dev, 115(1-2), 15–26.

Huang, M. C., Cheong, W. C., Lim, L. S., & Li, M. H. (2012). A simple, high sensitivity mutation screening using Ampligase mediated T7 endonuclease I and Surveyor nuclease with microfluidic capillary electrophoresis. Electrophoresis, 33(5), 788–796. doi:10.1002/elps.201100460

Itoh, M., Kim, C. H., Palardy, G., Oda, T., Jiang, Y. J., Maust, D., … Chitnis, A. B. (2003). Mind bomb is a ubiquitin ligase that is essential for efficient activation of Notch signaling by Delta. Dev Cell, 4(1), 67–82.

Jarriault, S., Schwab, Y., & Greenwald, I. (2008). A Caenorhabditis elegans model for epithelial-neuronal transdifferentiation. Proc Natl Acad Sci U S A, 105(10), 3790–3795. doi:10.1073/pnas.0712159105

Kalluri, R., & Weinberg, R. A. (2009). The basics of epithelial-mesenchymal transition. J Clin Invest, 119(6), 1420–1428. doi:10.1172/jci39104

Kimmel, C. B., Ballard, W. W., Kimmel, S. R., Ullmann, B., & Schilling, T. F. (1995). Stages of embryonic development of the zebrafish. Dev Dyn, 203(3), 253–310. doi:10.1002/aja.1002030302

Koo, B. K., Lim, H. S., Song, R., Yoon, M. J., Yoon, K. J., Moon, J. S., … Kong, Y. Y. (2005). Mind bomb 1 is essential for generating functional Notch ligands to activate Notch. Development, 132(15), 3459–3470. doi:10.1242/dev.01922

Kovacs, M., Toth, J., Hetenyi, C., Malnasi-Csizmadia, A., & Sellers, J. R. (2004). Mechanism of blebbistatin inhibition of myosin II. J Biol Chem, 279(34), 35557–35563. doi:10.1074/jbc.M405319200

Ku, H. Y., & Sun, Y. H. (2017). Notch-dependent epithelial fold determines boundary formation between developmental fields in the Drosophila antenna. PLoS Genet, 13(7), e1006898. doi:10.1371/journal.pgen.1006898

Lafkas, D., Shelton, A., Chiu, C., de Leon Boenig, G., Chen, Y., Stawicki, S. S., … Siebel, C. W. (2015). Therapeutic antibodies reveal Notch control of transdifferentiation in the adult lung. Nature, 528(7580), 127–131. doi:10.1038/nature15715

Lewis, W. H. (1947). Mechanics of invagination. Anat Rec, 97(2), 139–156.

Liu, Y., Pathak, N., Kramer-Zucker, A., & Drummond, I. A. (2007). Notch signaling controls the differentiation of transporting epithelia and multiciliated cells in the zebrafish pronephros. Development, 134(6), 1111–1122. doi:10.1242/dev.02806

Locy WA, L. O. (1916). The embryology of the bird’s lung based on observations of the bronchial tree. Part I. Am J Anat., 19, 447–504.

Ma, M., & Jiang, Y. J. (2007). Jagged2a-notch signaling mediates cell fate choice in the zebrafish pronephric duct. PLoS Genet, 3(1), e18. doi:10.1371/journal.pgen.0030018

Maddodi, N., & Setaluri, V. (2010). Prognostic significance of melanoma differentiation and trans-differentiation. Cancers (Basel), 2(2), 989–999. doi:10.3390/cancers2020989

Majumdar, A., Lun, K., Brand, M., & Drummond, I. A. (2000). Zebrafish no isthmus reveals a role for pax2.1 in tubule differentiation and patterning events in the pronephric primordia. Development, 127(10), 2089–2098.

Maki, N., Suetsugu-Maki, R., Tarui, H., Agata, K., Del Rio-Tsonis, K., & Tsonis, P. A. (2009). Expression of stem cell pluripotency factors during regeneration in newts. Dev Dyn, 238(6), 1613–1616. doi:10.1002/dvdy.21959

Maniotis, A. J., Folberg, R., Hess, A., Seftor, E. A., Gardner, L. M., Pe’er, J., … Hendrix, M. J. (1999). Vascular channel formation by human melanoma cells in vivo and in vitro: vasculogenic mimicry. Am J Pathol, 155(3), 739–752. doi:10.1016/s0002-9440(10)65173-5

Marra, A. N., & Wingert, R. A. (2016). Epithelial cell fate in the nephron tubule is mediated by the ETS transcription factors etv5a and etv4 during zebrafish kidney development. Dev Biol.

Martin, A. C., & Goldstein, B. (2014). Apical constriction: themes and variations on a cellular mechanism driving morphogenesis. Development, 141(10), 1987–1998. doi:10.1242/dev.102228

Martin, A. C., Kaschube, M., & Wieschaus, E. F. (2009). Pulsed actin-myosin network contractions drive apical constriction. Nature, 457(7228), 495. doi:10.1038/nature07522

Martin, A. C., Kaschube, M., & Wieschaus, E. F. (2009). Pulsed contractions of an actin-myosin network drive apical constriction. Nature, 457(7228), 495–499. doi:10.1038/nature07522

Mason, F. M., Tworoger, M., & Martin, A. C. (2013). Apical domain polarization localizes actin-myosin activity to drive ratchet-like apical constriction. Nat Cell Biol, 15(8), 926–936. doi:10.1038/ncb2796

Massa, F., Garbay, S., Bouvier, R., Sugitani, Y., Noda, T., Gubler, M. C., … Fischer, E. (2013). Hepatocyte nuclear factor 1beta controls nephron tubular development. Development, 140(4), 886–896. doi:10.1242/dev.086546

Menke, A. L., Spitsbergen, J. M., Wolterbeek, A. P., & Woutersen, R. A. (2011). Normal anatomy and histology of the adult zebrafish. Toxicol Pathol, 39(5), 759–775. doi:10.1177/0192623311409597

Merrell, A. J., & Stanger, B. Z. (2016). Adult cell plasticity in vivo: de-differentiation and transdifferentiation are back in style. Nat Rev Mol Cel Biol. doi:10.1038/nrm.2016.24

Moloney, D. J., Panin, V. M., Johnston, S. H., Chen, J., Shao, L., Wilson, R., … Vogt, T. F. (2000). Fringe is a glycosyltransferase that modifies Notch. Nature, 406(6794), 369–375. doi:10.1038/35019000

Naylor, R. W., & Davidson, A. J. (2014). Hnf1beta and nephron segmentation. Pediatr Nephrol, 29(4), 659–664. doi:10.1007/s00467-013-2662-x

Naylor, R. W., & Davidson, A. J. (2016). Pronephric tubule formation in zebrafish: morphogenesis and migration. Pediatr Nephrol. doi:10.1007/s00467-016-3353-1

Naylor, R. W., Przepiorski, A., Ren, Q., Yu, J., & Davidson, A. J. (2013). HNF1beta is essential for nephron segmentation during nephrogenesis. J Am Soc Nephrol, 24(1), 77–87. doi:10.1681/asn.2012070756

Naylor, R. W., Skvarca, L. B., Thisse, C., Thisse, B., Hukriede, N. A., & Davidson, A. J. (2016). BMP and retinoic acid regulate anterior-posterior patterning of the non-axial mesoderm across the dorsal-ventral axis. Nat Commun, 7, 12197. doi:10.1038/ncomms12197

Nikolopoulou, E., Galea, G. L., Rolo, A., Greene, N. D., & Copp, A. J. (2017). Neural tube closure: cellular, molecular and biomechanical mechanisms. Development, 144(4), 552–566. doi:10.1242/dev.145904

Panin, V. M., Papayannopoulos, V., Wilson, R., & Irvine, K. D. (1997). Fringe modulates Notch-ligand interactions. Nature, 387(6636), 908–912. doi:10.1038/43191

Qiu, X., Xu, H., Haddon, C., Lewis, J., & Jiang, Y. J. (2004). Sequence and embryonic expression of three zebrafish fringe genes: lunatic fringe, radical fringe, and manic fringe. Dev Dyn, 231(3), 621–630. doi:10.1002/dvdy.20155

Ray, H. J., & Niswander, L. A. (2016). Grainyhead-like 2 downstream targets act to suppress epithelial-to-mesenchymal transition during neural tube closure. Development, 143(7), 1192–1204. doi:10.1242/dev.129825

Sadler, T. W. (2005). Embryology of neural tube development. Am J Med Genet C Semin Med Genet, 135c(1), 2–8. doi:10.1002/ajmg.c.30049

Sammut, M., Cook, S. J., Nguyen, K. C., Felton, T., Hall, D. H., Emmons, S. W., … Barrios, A. (2015). Glia-derived neurons are required for sex-specific learning in C. elegans. Nature, 526(7573), 385–390. doi:10.1038/nature 15700

Sanchez Alvarado, A., & Tsonis, P. A. (2006). Bridging the regeneration gap: genetic insights from diverse animal models. Nat Rev Genet, 7(11), 873–884. doi:10.1038/nrg1923

Sawyer, J. M., Harrell, J. R., Shemer, G., Sullivan-Brown, J., Roh-Johnson, M., & Goldstein, B. (2010). Apical constriction: a cell shape change that can drive morphogenesis. Dev Biol, 341(1), 5–19. doi:10.1016/j.ydbio.2009.09.009

Scarlett, K., Pattabiraman, V., Barnett, P., Liu, D., & Anderson, L. M. (2015). The proangiogenic effect of iroquois homeobox transcription factor Irx3 in human microvascular endothelial cells. J Biol Chem, 290(10), 6303–6315. doi:10.1074/jbc.M114.601146

Schier, A. F., Neuhauss, S. C., Harvey, M., Malicki, J., Solnica-Krezel, L., Stainier, D. Y., … Driever, W. (1996). Mutations affecting the development of the embryonic zebrafish brain. Development, 123, 165–178.

Shekhani, M. T., Jayanthy, A. S., Maddodi, N., & Setaluri, V. (2013). Cancer stem cells and tumor transdifferentiation: implications for novel therapeutic strategies. Am J Stem Cells, 2(1), 52–61.

Shih, H. P., Kopp, J. L., Sandhu, M., Dubois, C. L., Seymour, P. A., Grapin-Botton, A., & Sander, M. (2012). A Notch-dependent molecular circuitry initiates pancreatic endocrine and ductal cell differentiation. Development, 139(14), 2488–2499. doi:10.1242/dev.078634

Stone, L. S. (1967). An investigation recording all salamanders which can and cannot regenerate a lens from the dorsal iris. J Exp Zool, 164(1), 87–103. doi:10.1002/jez.1401640109

Tapscott, S. J., Davis, R. L., Thayer, M. J., Cheng, P. F., Weintraub, H., & Lassar, A.B. (1988). MyoD1: a nuclear phosphoprotein requiring a Myc homology region to convert fibroblasts to myoblasts. Science, 242(4877), 405–411.

Thisse, C., & Thisse, B. (2008). High-resolution in situ hybridization to whole-mount zebrafish embryos. Nat Protoc, 3(1), 59–69. doi:10.1038/nprot.2007.514

Thorel, F., Nepote, V., Avril, I., Kohno, K., Desgraz, R., Chera, S., & Herrera, P. L. (2010). Conversion of adult pancreatic alpha-cells to beta-cells after extreme beta-cell loss. Nature, 464(7292), 1149–1154. doi:10.1038/nature08894

Thowfeequ, S., Myatt, E. J., & Tosh, D. (2007). Transdifferentiation in developmental biology, disease, and in therapy. Dev Dyn, 236(12), 3208–3217. doi:10.1002/dvdy.21336

Tosh, D., & Slack, J. M. (2002). How cells change their phenotype. Nat Rev Mol Cell Biol, 3(3), 187–194. doi:10.1038/nrm761

Tseng, D. Y., Chou, M. Y., Tseng, Y. C., Hsiao, C. D., Huang, C. J., Kaneko, T., & Hwang, P. P. (2009). Effects of stanniocalcin 1 on calcium uptake in zebrafish (Danio rerio) embryo. Am J Physiol Regul Integr Comp Physiol, 296(3), R549–557. doi:10.1152/ajpregu.90742.2008

Ullrich, K. J., & Murer, H. (1982). Sulphate and phosphate transport in the renal proximal tubule. Philos Trans R Soc Lond B Biol Sci, 299(1097), 549–558.

Vasilyev, A., Liu, Y., Mudumana, S., Mangos, S., Lam, P. Y., Majumdar, A., … Drummond, I. A. (2009). Collective cell migration drives morphogenesis of the kidney nephron. PLoS Biol, 7(1), e9. doi:10.1371/journal.pbio.1000009

Verdeguer, F., Le Corre, S., Fischer, E., Callens, C., Garbay, S., Doyen, A., … Pontoglio, M. (2010). A mitotic transcriptional switch in polycystic kidney disease. Nat Med, 16(1), 106–110. doi:10.1038/nm.2068

Wingert, R. A., & Davidson, A. J. (2008). The zebrafish pronephros: a model to study nephron segmentation. Kidney Int, 73(10), 1120–1127. doi:10.1038/ki.2008.37

Wingert, R. A., & Davidson, A. J. (2011). Zebrafish nephrogenesis involves dynamic spatiotemporal expression changes in renal progenitors and essential signals from retinoic acid and irx3b. Dev Dyn, 240(8), 2011–2027. doi:10.1002/dvdy.22691

Wingert, R. A., Selleck, R., Yu, J., Song, H.-D., Chen, Z., Song, A., … McMahon, A. P. (2007). The cdx genes and retinoic acid control the positioning and segmentation of the zebrafish pronephros. PLoS Genet, 3(10), 1922–1938.

Yanger, K., Zong, Y., Maggs, L. R., Shapira, S. N., Maddipati, R., Aiello, N. M., … Stanger, B. Z. (2013). Robust cellular reprogramming occurs spontaneously during liver regeneration. Genes Dev, 27(7), 719–724. doi:10.1101/gad.207803.112

Ye, L., Robertson, M. A., Hesselson, D., Stainier, D. Y., & Anderson, R. M. (2015). Glucagon is essential for alpha cell transdifferentiation and beta cell neogenesis. Development, 142(8), 1407–1417. doi:10.1242/dev.117911

